# Corticosterone differentially modulates time-dependent fear generalization following mild or moderate fear conditioning training in rats

**DOI:** 10.1101/2021.03.10.434749

**Authors:** Moisés dos Santos Corrêa, Barbara dos Santos Vaz, Beatriz Scazufca Menezes, Tatiana Lima Ferreira, Paula Ayako Tiba, Raquel Vecchio Fornari

## Abstract

Stressful and emotionally arousing experiences create strong memories that seem to lose specificity over time. It is uncertain, however, how the stress system contributes to the phenomenon of time-dependent fear generalization. Here, we investigated whether post-training corticosterone (CORT-HBC) injections, given after different training intensities, affect contextual fear memory specificity at several time points. We trained male Wistar rats on the contextual fear conditioning (CFC) task using two footshock intensities (mild CFC, 3 footshocks of 0.3 mA, or moderate CFC, 3x 0.6 mA) and immediately after the training session we administered CORT-HBC systemically. We first tested the animals in a novel context and then in the training context at different intervals following training (2, 14, 28 or 42 days). By measuring freezing in the novel context and then contrasting freezing times shown in both contexts, we inferred contextual fear generalization for each rat, classifying them into Generalizers or Discriminators. Following mild CFC training, the glucocorticoid injection promoted an accurate contextual memory at the recent time point (2 days), and increase the contextual memory accuracy 28 days after training. In contrast, after the moderate CFC training, CORT-HBC facilitated contextual generalization at 14 days, compared to the control group that maintained contextual discrimination at this timepoint. For this training intensity, however, CORT-HBC did not have any effect on recent memory specificity. These findings indicate that treatment with CORT-HBC immediately after the encoding of mild or moderately arousing experiences may differentially modulate memory consolidation and time-dependent fear generalization.

**HIGHLIGHTS:**

- Rats were trained in a contextual fear task and tested for fear generalization.
- Corticosterone (CORT) injected after a mild training promoted accuracy up to 28 days.
- CORT injected after a moderate training facilitated time-dependent fear generalization.

## INTRODUCTION

Stressful events elicit the formation of lasting and intense memories and their effects on memory is thought to be exerted by temporally contiguous activation of the noradrenergic and glucocorticoid systems (Joëls et al., 2011; Vogel et al., 2016). Even though these two hormonal systems have been long associated with the modulation of the strength of fear memories (de Quervain et al., 2016; Roozendaal et al., 2009), their role on memory specificity has only recently become a new focus of investigation. Moreover, overgeneralization of fear is associated with anxious and phobic behavior and is one of the main symptoms of several stress-related disorders, such as post-traumatic stress disorder (PTSD), generalized anxiety disorder and phobias (Lenaert et al., 2015; Lissek et al., 2014b, 2014a). Considering the widespread use of corticosteroids for the treatment of a myriad of diseases (Rice et al., 2017), understanding how stress modifies not only memory strength but also memory specificity has deep implications in the prevention of these disorders and the development of better treatments for them.

A growing body of evidence suggests that noradrenaline and corticosterone (CORT) modulate memory acquisition and consolidation by shifting from cognitive detailed contextual learning towards a more rigid memory system, associated with habit and stimulus response memories (Schwabe, 2017). Such evidence suggests that generalized fear is associated with this rigid, habit-forming memory system, favoring rapid reactions toward threatening stimuli (Quaedflieg and Schwabe, 2018). Likewise, several other studies in humans found association between high levels of cortisol, during or immediately after memory acquisition, and a loss of contextual dependency in episodic memories (van Ast et al., 2013). Furthermore, Roozendaal and Mirone have found, in rats, that acute CORT treatment after training in a modified version of the inhibitory avoidance task elicited higher fear generalization in episodic-like memories (Roozendaal and Mirone, 2020). Accumulated evidence have shown that, in fact, high plasma CORT levels after an aversive situation increase the interaction between the basolateral nucleus of the amygdala and the prefrontal cortex, eliciting the consolidation of a more ‘generalizable’ memory trace (Bahtiyar et al., 2020). On the other hand, in humans, cortisol seemed to impair generalization in low-arousing types of associative learning (Dandolo and Schwabe, 2016) and enhance contextualization of emotionally charged pictures (Van Ast et al., 2014). Hence, it is possible that the glucocorticoid system modifies the specificity of emotionally charged memories, but the orientation of this modulation is still controversial.

Parallel to the investigation of how stress modulates recent memory generalization, decades of research have shown that the process of systems consolidation -- in which the retrieval of contextual memory traces supposedly become more dependent of neocortical regions over time -- induces a loss of contextual details in episodic memories with the passage of time (Moscovitch et al., 2016). This loss of contextual specificity is associated with time-dependent fear generalization across different contexts (Tonegawa et al., 2018). Even though there is evidence associating stress with generalized contextual memories, few studies investigated whether stress or high arousal levels during training alter systems consolidation and the phenomenon of time-dependent fear generalization (Atucha et al., 2017; Bahtiyar et al., 2020; Pedraza et al., 2016).

To elucidate this issue, a recent study showed that a mildly arousing contextual fear conditioning (CFC) training in rats (with 3 footshocks of 0.3 mA) induced low levels of post-training plasma CORT and persistent contextual discrimination at 14 and 28 days after training. On the other hand, moderate CFC training (3x 0.6 mA footshocks) elicited higher levels of post-training plasma CORT and a generalized contextual fear response 28 days later. Interestingly, animals trained with moderate CFC and tested 14 days later showed a generalized fear memory in the first experiment, and a discriminative fear response in the second (replication) experiment. In addition, freezing behavior of rats tested in the novel context at this time point was linearly associated with individual post-training plasma CORT levels, suggesting that there could be an interaction between training arousal level and post-training glucocorticoid activation that facilitates fear generalization, depending on individual variability (dos Santos Corrêa et al., 2019). This evidence suggests that systems consolidation is a complex process that expresses itself differently after mild or highly arousing CFC, but it also raises the question of whether this process can be further tempered by artificially activating the glucocorticoid system via systemic treatment with CORT. The present study, therefore, investigated the modulation of time-dependent fear generalization exerted by CORT administration following mild and moderate CFC trainings.

## METHODS

### 2.1 Subjects

Three-month old male Wistar rats, obtained from *Centro de Desenvolvimento de Modelos Experimentais para Biologia e Medicina* (CEDEME-UNIFESP, total n = 224; weighing between 275-385g at time of training), were kept in controlled conditions of temperature (23 ± 2°C), humidity and light/darkness cycle of 12:12 hours (light phase starting at 7am). Rats were randomly housed in groups of 4 per home-cage (30 cm × 16 cm × 28 cm) and provided with food and water *ad libitum*. Wooden flakes were used as bedding in the cages. The rats were adapted to the vivarium for at least one week before the beginning of the experiments. A paper towel in each cage was used as environmental enrichment. Sanitation was provided twice a week by the vivarium technicians. Training and testing were performed during the light phase of the cycle (between 10:00am-03:00pm), at the rat’s nadir of the diurnal rhythm for corticosterone. The principal experimenter was a male. All procedures were conducted according to the guidelines and standards of CONCEA - *Conselho Nacional de Controle de Experimentação Animal* (Brazilian Council of Animal Experimentation) and were previously approved by the Ethics Committee on Animal Use - UFABC (CEUA - protocol numbers 5553080618 and 7734310719).

### 2.2 Systemic Drug administration

A preformed water-soluble complex of corticosterone and 2-hydroxypropyl-β-cyclodextrin (CORT-HBC, 0.25, 0.5, 1.0, 2.0, 4.0 or 8.0 mg/kg, C174 Sigma-Aldrich, Germany) dissolved in sterile saline, in a volume of 2 ml/kg body weight, was administered subcutaneously immediately after the training session. We chose the drug, solubility and other parameters based on previous studies (Kaouane et al., 2012; Rusu et al., 2018; Timmer and Sandi, 2010). Drug administration took place in the training room, opposite from the conditioning chambers. Doses of CORT-HBC were selected based on a previous study that showed an effect on contextual fear memory in mice with this specific molecular compound (Kaouane et al., 2012) and a standardization experiment was done to choose the best doses for our experimental design with Wistar rats. Drug solutions were freshly prepared before each experiment.

### 2.3 Apparatus

Behavioral experiments were conducted in two identical automated fear conditioning chambers (Med-Associates, Inc., St. Albans, VT) connected to a computer interface enabling video recording, analysis, and automated measurement of the rat’s freezing behavior in real time. The chambers and software used were previously described in detail (de Paiva et al., 2021; dos Santos Corrêa et al., 2019). Freezing measure was automated (VideoFreeze Version 1.12.0.0, MedAssociates) and set for less than 20 movement units for at least 30 frames (1s) (Bueno et al., 2017; Moreira-Silva et al., 2018).

The training context (context A) was characterized by a grid floor composed of 20 stainless steel rods (diameter: 4.8mm), top and front walls made of transparent polycarbonate, a back wall made of white acrylic covered with a 5-cubed white-on-black pattern, stainless-steel sidewalls, and a drop pan below the grid floor. The light in the conditioning box remained on and only the exhaust fan noise was emitted during the training and test sessions. Context A was cleaned with alcohol 10% before and after each session.

The novel context (Context B) consisted of the conditioning chamber and stainless-steel drop pan personalized with a grid floor (20 interleaved rods of either 4.8 or 9.5mm of diameter) and white Plexiglas curved sidewalls extending across the back wall. The light in the box remained turned off and a 90-dB white noise was emitted in addition to the fan noise during the test. Context B was cleaned with a 5% acetic acid solution before and after each session.

### 2.4. Contextual Fear Conditioning task and experimental designs

In all experiments, rats from each home-cage were randomly assigned to one of the experimental or control groups, and all groups were matched according to the average body weight. Rats were handled for 3 days prior to the CFC training, each for 3 min, in the training room. They were habituated to an adjacent room for 40 to 90 minutes before the handling sessions and both training and testing. To minimize potential confounders, the same experimenter always handled the rats.

On the training session, rats were individually transported by the experimenter from the adjacent room to the training room in a white acrylic, individual cage (to which they were accustomed during habituation days) and placed in context A. After 2 minutes of free exploration, three footshocks (0.3 or 0.6 mA, 1s each, depending on the experiment) were delivered with intervals of 30s between them. One minute after the third shock, the animal was removed from the apparatus and immediately received a subcutaneous (s.c) injection of saline or different doses of CORT-HBC. The same experimenter that transported the animals to the training chamber removed them from the apparatus and administered the injections. Each animal was then returned to its individual cage and transported back to the adjoining room where it stayed for at least 90 minutes before being placed back in it home-cage and returned to the vivarium.

For test sessions, rats were transported from the adjacent room to the training room in a wooden box (which was novel to the animals), by a different experimenter from the training session, unaware of the animal’s treatment group, and placed in context B for 4 minutes. Afterwards, each animal was removed from this conditioning chamber by the principal experimenter, placed into the white acrylic individual cage for up to 1 minute, and then exposed to context A, for 4 minutes. In both contexts, no shock was delivered during the test and animal’s freezing time was automatically measured by the software. Both the blindness of the different experimenter and the automated freezing measurement reduced the potential risk of bias. Figure 1 shows an overview of the experimental procedures.

**Figure 1:**
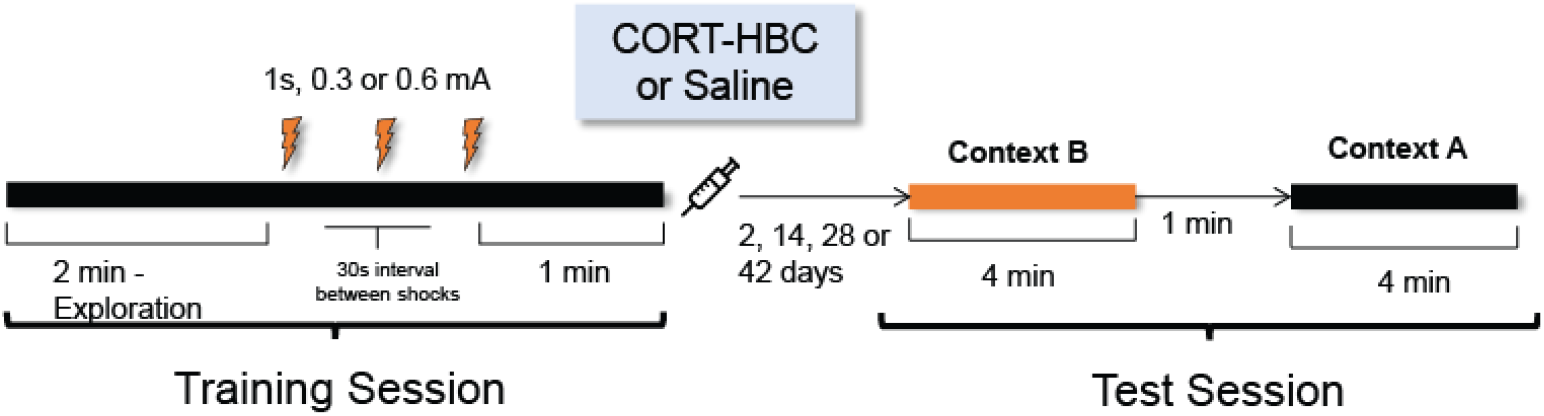
Schematic representation of the experimental designs. Rats were exposed to the training context for two minutes, when they were free to explore. Thereafter, they received three footshocks of one intensity, 0.3 or 0.6mA (1s each, 30s between them). Following the last footshock, each rat stayed in the chamber for another minute and was then removed. Immediately after training, the rats received an injection of CORT-HBC or saline, according to their experimental group. After 2, 14, 28 or 42 days, rats were tested for long-term memory. They were first exposed to a novel context (Context B) for 4 minutes and then returned to their individual cage, where they stayed for 1 minute. Then, they were exposed to the training context (Context A) for 4 minutes.

#### 2.4.1 Dose selection experiment

A dose-response curve was performed to verify the best doses of CORT-HBC for our experimental design with Wistar rats. Rats were trained with three footshocks (0.3 mA) and, immediately after training, received a s.c injection of saline or CORT-HBC (0.25, 0.5, 1.0, 2.0, 4.0 or 8.0 mg/kg). The test sessions were performed 2 days after training, as described above.

#### 2.4.2 Corticosterone modulation of recent and remote contextual fear memory specificity

In order to investigate whether the treatment with CORT-HBC modulates contextual fear specificity over time following mild or moderate CFC training, different testing timepoints were chosen based on the study of dos Santos Corrêa et al. (2019). In this previous study, mild CFC (0.3mA) elicited discriminative fear memory up to 28 days, whereas moderate CFC (0.6mA) resulted in contextual generalization or discrimination 14 days after training, in two different experiments. These divergent results on the 14^th^ day may be explained by the different levels of post-training plasma CORT observed between both groups of rats (see Figure S1 in the supplementary material). For the present study, we hypothesized that injections of CORT-HBC immediately after a mild CFC training would promote fear generalization at remote timepoints (28 or 42 days after training), whereas the same treatment after a moderate CFC training would facilitate fear generalization at the 14th day timepoint. Therefore, rats were subject to the CFC training with one of two shock intensities (3x of 0.3 or 0.6mA) and immediately after training received a s.c. injection of saline or CORT-HBC (4.0 or 8.0 mg/kg). Rats trained with 0.3mA footshocks were tested 28 or 42 days after training, whereas those trained with the 0.6mA intensity were tested 2 or 14 days after training. Different groups of rats were used for each footshock intensity and timepoint.

### 2.5 Statistical analysis

For each experiment, the experimental unit was the individual rat. All descriptive and inferential statistical analysis was done using JAMOVI (version 1.63). Fear response to footshocks was quantified as the time the animal spent in freezing during the last minute of the training session. The conditioned fear response to the training context was quantified as the time the animal spent in freezing in context A, whereas fear generalization was considered as the time of freezing in context B during the recall tests. Behavioral results are expressed as the group mean percent freezing time ± standard error of the mean (S.E.M). An “index of generalization” was calculated using the formula 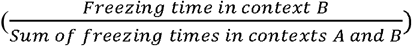 and are expressed as group mean generalization (± S.E.M). A generalization threshold of 0.3 was set to distinguish rats between ‘discriminator’ (< 0.30) and ‘generalized animals (≥ 0.30), a level that is equivalent to the one reported by Wiltgen and colleagues (Wiltgen et al., 2010). When needed to satisfy the requirements for the use of the ANOVA, total freezing times were transformed using the square root transform function (dos Santos Corrêa et al., 2019) to make data normally distributed and homogeneous, according to the tests of Kolmogorov-Smirnov and Levene. When not stated otherwise, data used in the inferential tests was normally distributed and homogenous. All statistical analysis are stored together with the behavioral data at https://data.mendeley.com/datasets/7xvshh2s59/.

Freezing data from the last minute of the training sessions (post-shock freezing times) were analyzed with one-way ANOVAs with Treatment as the independent factor. Freezing data from the test sessions were analyzed with one-way or repeated measures ANOVAs (RMANOVAs, with freezing time in Contexts as within-subject dependent variables and Treatment as an independent factor). Generalization index data were analyzed with one factor ANOVAs with Treatment as the independent factor. The Tukey *post hoc* test was further used to identify significant differences when applicable. Bonferroni corrections were used for multiple comparisons when needed. For the non-homogenous Generalization Index data, the Welch corrected ANOVA was used to analyze means and the Games-Howell Tukey corrected post hoc tests were used to compare differences between groups. The tests ‘linear tendency of Cochran-Armitage’ (experiment 1 only) and ‘likelihood-ratio chi-square’ were used to test the difference in the distribution of ‘generalized and ‘discriminator’ animals across all treatment groups.

Significance for all inferential tests was set at *p* ≤ 0.05. Effect sizes for ANOVAs and RMANOVAs (*η^2^p* or *η^2^G*, respectively) are reported only when the test was found significant (“*η^2^p*” or *“η^2^G*” values above 0.14 are considered large effects; values between 0.06 and 0.14 are considered moderate; and below 0.06, small). For post hoc tests, the Cohen’s d value is reported as a measure of effect size when the test was found significant. All four measures (statistical difference in mean freezing times in the novel context, lack of difference between the training and the novel contexts, difference in mean generalization index and difference in proportion of generalizers per group) were used to consider group level generalization. Significant results in at least two of these measures were considered as an evidence of fear generalization in any particular group.

## RESULTS

### 3.1 CORT-HBC dosage selection

In order to verify and select which CORT-HBC doses would be effective in modulating contextual fear memory consolidation, different groups of rats received a subcutaneous injection of saline or CORT-HBC (0.25, 0.5, 1.0, 2.0, 4,0 and 8.0 mg/kg) immediately after training and were tested 2 days later as described above. Results indicate that higher doses of CORT-HBC seem to promote recent memory accuracy (Figure 2). The RMANOVA showed a significant Context effect [*F* _(1, 40)_ = 39.73, *p* < 0.01, *η^2^G* = 0.24], but no Treatment effect [*F* _(6, 40)_ = 0.44, *p* = 0.85] or interaction effect between Context * Treatment [*F* _(6, 40)_ = 1.12, *p* = 0.37, Fig. 2a]. The same result was confirmed by analyzing the mean Generalization Index [*F* _(6, 40)_ = 1.58, *p* = 0.18, Fig. 2b]. Nonetheless, a trend toward a differential distribution in the number of animals that showed Generalization Indices lower than 0.3 (Discriminators), as the dose of CORT-HBC increased, was confirmed by the test of linear tendency of Cochran-Armitage [χ^2^ _(1, 47)_ = 9.4, *p* < 0.01] and the likelihood-ratio chi-square test χ^2^ _(6, 47)_ = 12.68, *p* = 0.048, Fig. 2c].

**Figure 2:**
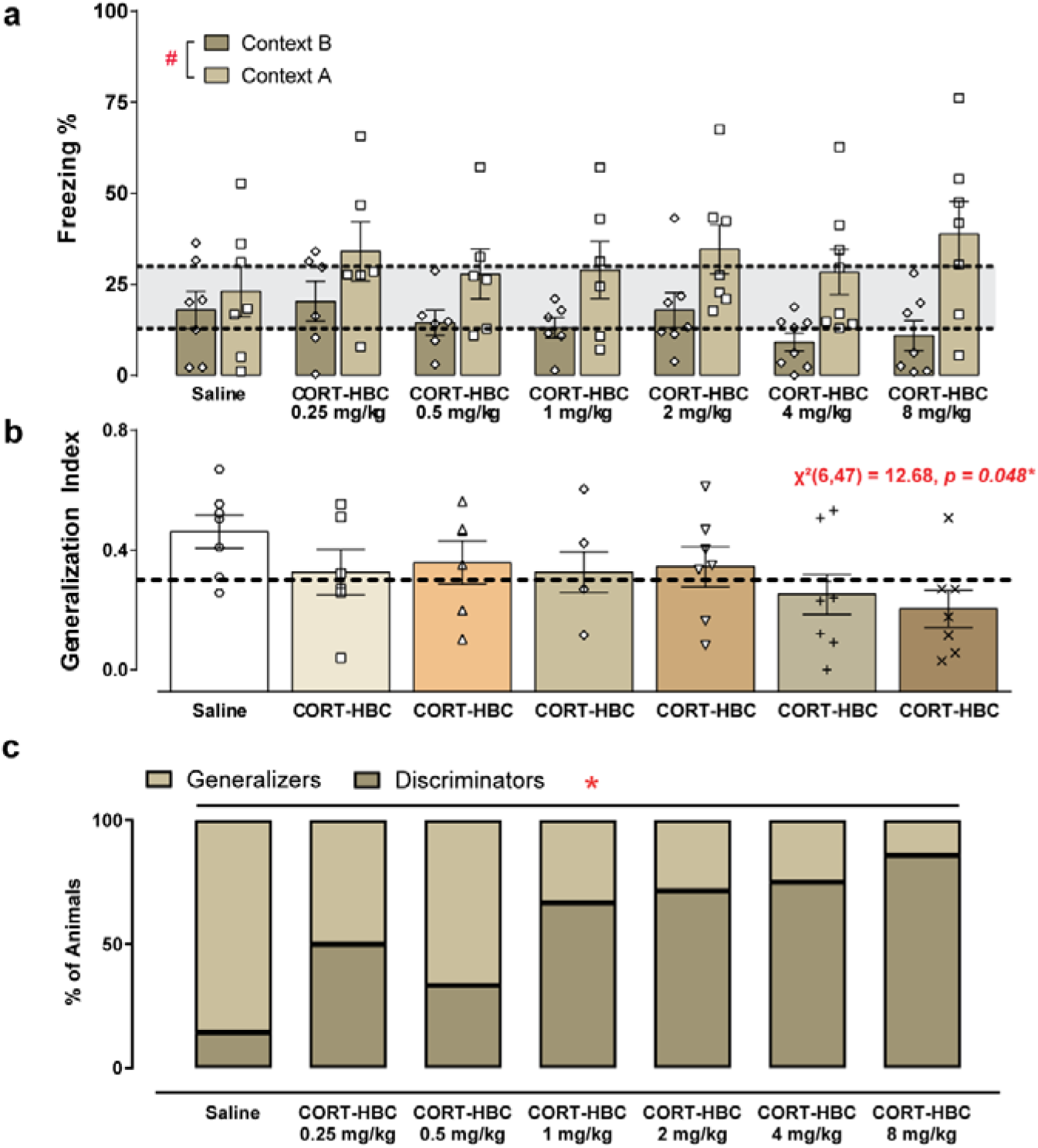
Post-training corticosterone-HBC (CORT-HBC) modulates recent (2 days) memory specificity. (a) Mean freezing percentage ± standard error of the mean (S.E.M) shown in the novel context (B, diamond symbol) and then in the training context (A, square symbol) of groups injected with Saline (n = 7), CORT-HBC 0.25 mg/kg (6), 0.5 mg/kg (6), 1 mg/kg (6), 2 mg/kg (7), 4 mg/kg (7) and 8 mg/kg (7), immediately after mild CFC training (3 footshocks of 0.3 mA). The dotted lines and gray area represent the lower error bar in context B and the higher error bar in context A for the saline group. (#) p < 0.01 in the main effect of the factor Context. (b) Mean (± S.E.M.) Generalization Index [B/(A+B)] of each treatment group. The dotted line represents the threshold point between discriminators (0 – 0.3) and generalizers (0.31-1). (c) Percentage of animals that were considered “discriminators” (below) or “generalizers” (above). The percentage of discriminators shows a linear increase in response to increasing doses of CORT-HBC (*) p < 0.01 in the Cochran-Armitage test for linear association).

### 3.2 Corticosterone differentially modulates fear memory specificity for mild and moderate CFC training intensities over time

#### 3.2.1 Mild CFC training intensity

To allow comparison of the treatment effect found above with all the other experiments, a subset of the data presented at 3.1 was used to further analyze fear generalization of rats treated with the highest doses of CORT-HBC (4.0 or 8.0 mg/kg, s.c.) or saline, immediately after the mild intensity CFC training, and tested 2 days later (Fig. 3a-c). There was no difference in post-shock freezing levels between these groups during training (Treatment [*F* _(2, 19)_ = 0.09, *p* = 0.92], Table S1). For the test session, CORT-HBC treatment increased the difference in rat’s freezing percentage between contexts B and A in the recent memory time point. RMANOVA indicated a significant effect for Context [*F* _(1, 19)_ = 26.96, *p* < 0.001, *η^2^G* =0.26] and for the interaction Context * Treatment [*F* _(2, 19)_ = 3.82, *p* = 0.04, *η^2^G* = 0.09], but not for Treatment [*F* _(2, 19)_ = 0.38, *p* = 0.69]. Interaction post hoc tests confirmed that rats treated with Saline expressed similar levels of freezing in both contexts (*t* _(30.11)_ = −0.85, *p* = 0.95) whereas CORT-HBC treated animals showed lower freezing times when exposed to context B than when exposed to context A (CORT 4.0mg/kg, *t* _(30.11)_ = −3.46, *p* = 0.03, *d* = −1,60; CORT 8.0mg/kg, *t* _(30.11)_ = −4.72, *p* = 0.002, *d*=−2.16, Fig. 3a). For the Generalization Index, the one-way ANOVA showed a significant Treatment effect [*F* _(2, 19)_ = 4.67, *p* = 0.02, *η^2^p* =0.33] and the post hoc tests revealed that the 8.0 mg/kg-CORT-HBC group showed lower generalization levels than the Saline group (*t* _(19)_ = −2.86, *p* = 0.03, *d* =−1.31). Other comparisons were not statistically significant (*p* > 0.07, Fig. 3b). The likelihood-ratio chi-square test has further confirmed a difference in the percentage of discriminators’ rats in CORT-HBC treated groups (N=6 discriminators for both 4,0mg/kg and 8.0mg/kg doses) when compared to the saline-treated animals [N = 1 discriminator, χ^2^_(2, 22)_ = 9.29, p = 0.01, Fig. 3c].

**Figure 3:**
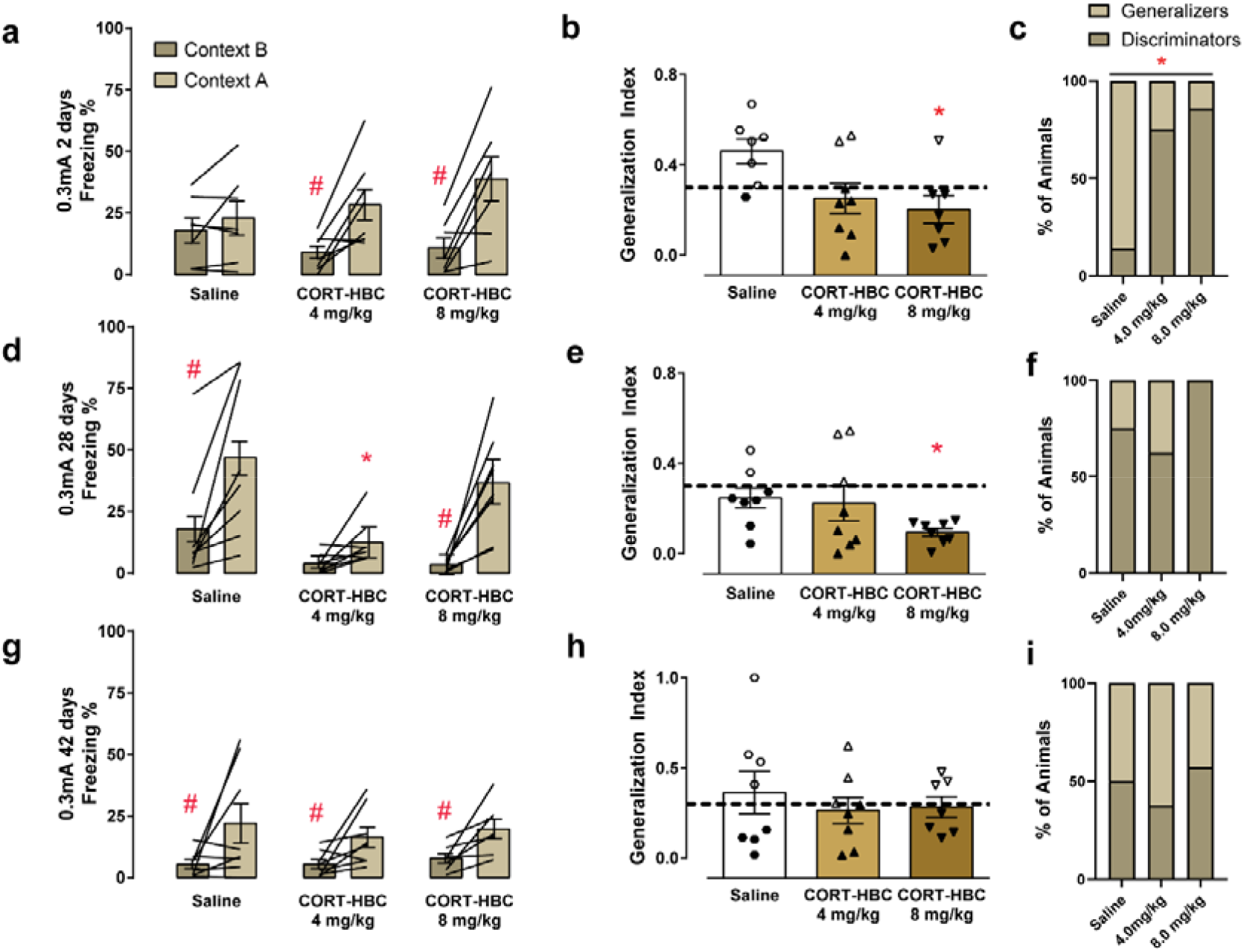
Corticosterone administration after mild intensity CFC training increases contextual memory specificity up to 28 days. (a, d, g) Mean freezing percentage ± Standard error of the mean (S.E.M) shown by rats exposed to the novel context (B) and then to the training context (A) after (a) 2, (d) 28 or (g) 42 days after training. Lines represent individual freezing data for each rat in context B and then in A. (#) p ≤ 0.001 when compared to within-subject freezing expressed in Context A. (*) p = 0.02 when compared to the mean freezing times expressed by the Saline group in Context A. (b, e, h) Mean Generalization Index ± S.E.M. [B/(A+B)] after (b) 2, (e) 28 or (h) 42 days after CFC. The dotted line represents the threshold between “discriminators” and “generalizers”. Filled symbols represent the animals considered discriminators and the unfilled symbols represent the ones considered generalizers. (*) p < 0.05 when compared to the Saline group. (c, f, i) Percentage of generalizer and discriminator animals per group after (c) 2, (f) 28 or (i) 42 days after CFC. (*) p < 0.05 in the proportion test. Number of rats for each group: 2 days, Saline (N = 7), CORT-HBC 4 mg/kg (N = 8), 8 mg/kg (N = 7). 28 days, Saline (N = 8), CORT-HBC 4 mg/kg (N = 8), 8 mg/kg (N = 8). 42 days, Saline (N = 8), CORT-HBC 4 mg/kg (N = 8), 8 mg/kg (N = 7).

Following the recent memory test, we chose two time points *a priori*, based on the results found by dos Santos Corrêa and colleagues (2019), to test our hypothesis that CORT-HBC would elicit generalized fear at remote timepoints. Two additional sets of rats were trained with the mild CFC intensity (0.3 mA), received a s.c. injection of CORT-HBC (4.0 or 8.0 mg/kg) or saline immediately after training and were tested 28 or 42 days later. For the first set of animals (that were tested at the 28^th^ day), there was no difference in post-shock freezing levels during training (Treatment [*F* _(2, 21)_ = 0.50, *p* = 0.61], Table S1). During the test session, contrary to our hypothesis, rats injected with the highest dose of CORT-HBC (8.0 mg/kg) showed a lower generalization across contexts, when compared to the control group (Fig. 3d-f). As freezing data in contexts B and A did not have homogenous variance (p < 0.05 for both), they were transformed using the square root transform and the RMANOVA was calculated using the transformed data. The RMANOVA revealed significant effects for Context [*F* _(1, 21)_ = 59.92, *p* < 0.001, *η^2^G* = 0.44] and Treatment [*F* _(2, 21)_ = 5.67, *p* = 0.01, *η^2^G* = 0.28]. The interaction between Context and Treatment showed a nearly significant effect [*F* _(2, 21)_ = 3.34, *p* = 0.055, *η^2^G* = 0.08], but as the generalized eta squared of 0.08 indicates a moderate effect size, unprotected post hoc tests were calculated. Post hoc tests revealed that groups injected with 8.0 mg/kg of CORT-HBC (*t* _(21)_ = −6.36, *p* < 0.001, *d*=−2.77) or saline (*t* _(21)_ = −4.35, *p* = 0.003, *d*=− 1.90) showed higher freezing levels in context A than in context B. In contrast, rats injected with 4.0 mg/kg of CORT-HBC displayed similar freezing levels in both contexts (*t* _(21)_ = −2.71, *p* = 0.12). Besides, rats in the 4.0 mg/kg-CORT-HBC group showed lower freezing times in context A when compared to the Saline group in the same context (*t* _(34.7)_ = −3.48, *p* = 0.02, *d*=−1.18). Other post hoc comparisons were not significant (*p* > 0.09). To further assess between-group differences, additional Bonferroni corrected ANOVAs were used to compare freezing in B and A separately (Table 1). These ANOVAs confirmed that rats in the 4.0 mg/kg-CORT-HBC group showed lower freezing times when compared to the Saline group in both contexts (*p* = 0.01 *and p* = *0.*047 in context A and B, respectively) whereas the 8.0 mg/kg-CORT-HBC group showed a non-significant trend towards a decrease in freezing percentage in the novel context, when compared to the saline group in this same context (*p* = 0.06, Table 1; for all other one-way ANOVAs see Table S2). For the Generalization Index, the Welch corrected ANOVA was used due to non-homogeneity of variance (*p* < 0.05). There was a significant Treatment effect [*F* _(2, 10.92)_ = 5.50, *p* = 0.02, *η^2^p* = 0.19], and the post hoc tests revealed that animals injected with 8.0 mg/kg of CORT-HBC presented lower generalization levels in comparison to the Saline group (*t* _(9.1)_ = −3.13, *p* = 0.03, *d*=−2.08), whereas the 4.0 mg/kg-CORT-HBC group showed a mean generalization index similar to the Saline (*t* _(11.3)_ = −0.26, *p* = 0.97) and to the 8.0 mg/Kg-CORT-HBC groups (*t* _(7.7)_ = 1.63, *p* = 0.29, Fig. 3e) and below the threshold of generalization. The likelihood-ratio chi-square test, however, did not show differences in the proportion of discriminators across all groups [χ^2^ _(2, 24)_ = 4.98, p = 0.08, Fig. 3f].

**Table 1:**
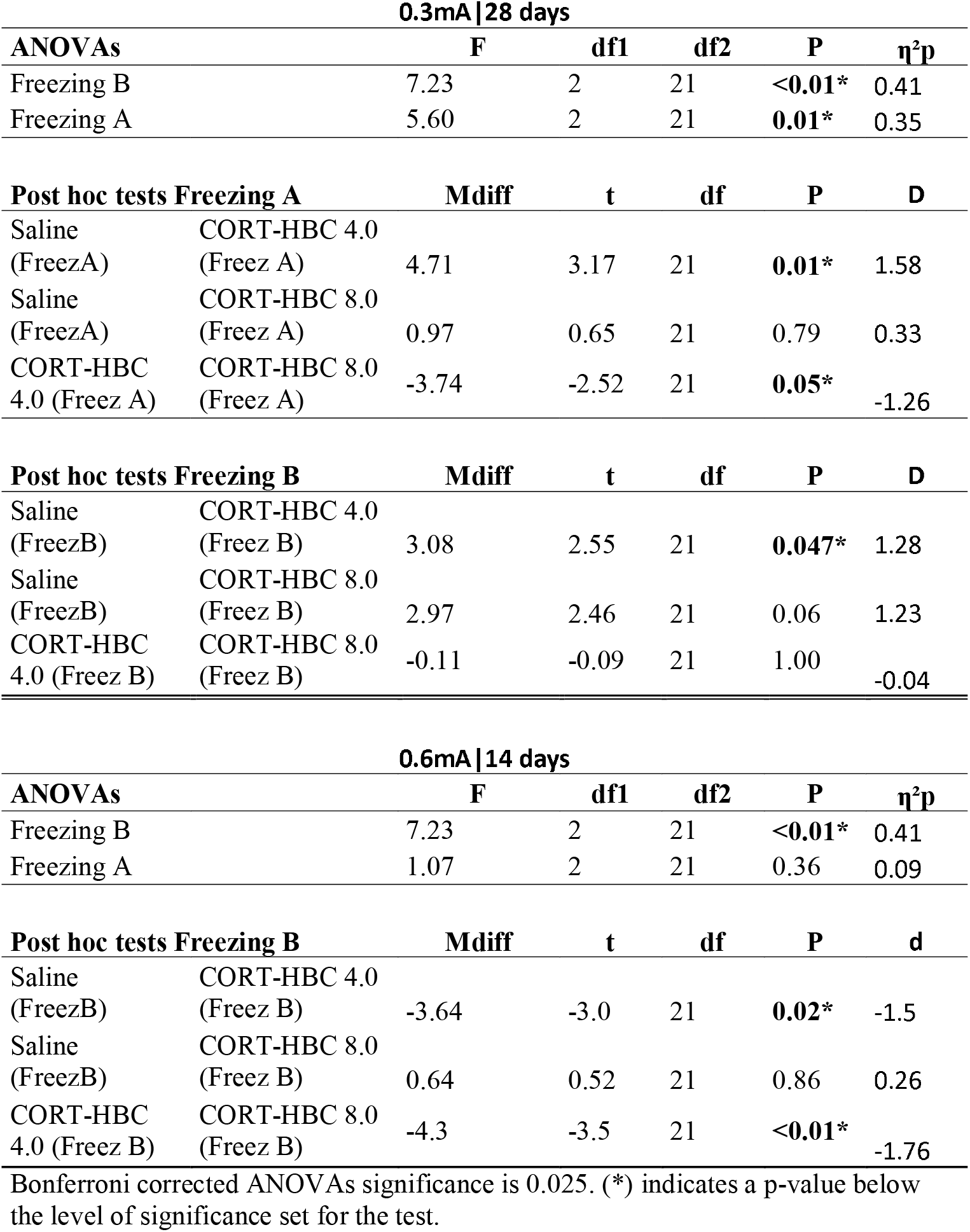
Bonferroni corrected One-way ANOVAs for the freezing times in the training (A) or novel (B) contexts and post hoc tests.

All rats tested on the 42nd day after training, on the other hand, showed higher freezing levels in Context A than in Context B and similar generalization levels (Fig. 3g-i). For this set of animals, there was no difference in post-shock freezing levels during training (Treatment [*F* _(2, 20)_ = 0.27, *p* = 0.77], Table S1). During the test session at this time point, freezing time data in context A was non-homogenous (*p* < 0.05) and therefore the transformed data was analyzed via RMANOVA. There was a significant effect for Context [*F* _(1, 20)_ = 13.93, *p* = 0.001, *η^2^G* = 0.26] but not for Treatment [*F* _(2, 20)_ = 0.29, *p* = 0.75] or the interaction between Context and Treatment [*F* _(2, 20)_ = 0.26, *p* = 0.77]. Mean generalization index was similar in all Treatment groups [*F* _(2, 20)_ = 0.37, *p* = 0.70, Fig. 3h] and this result was further confirmed by the likelihood-ratio chi-square test [χ^2^ _(2, 23)_ = 0.26, p = 0.88, Fig. 3i].

In summary, for the mild CFC training intensity, the post-training treatment with the highest dose of CORT-HBC (8.0 mg/kg) promoted contextual discrimination at 2 days and induced an enhancement of contextual memory accuracy at 28, but not 42, days after training. Additionally, rats injected with 4.0 mg/kg of CORT-HBC showed less freezing in both contexts at the 28^th^ day when compared to Saline-treated rats, but they preserved the generalization index under the threshold of generalization and this group showed a similar proportion of generalizer animals when compared to the other two groups.

#### 3.2.2 Moderate CFC training intensity

To verify whether CORT injections after a moderate CFC would facilitate or accelerate contextual fear generalization, different sets of rats were trained with 3 footshocks of 0.6mA, received s.c. injections of CORT-HBC (4.0 or 8.0 mg/kg) or saline immediately after training and were tested 2 or 14 days later. The results from the first set of animals, tested for the recent memory (2 days after training), indicate that all groups of rats showed higher freezing response in the training context than in the novel context, unabated by treatment (Fig. 4a). Again, there was no difference in post-shock freezing times during training (Treatment [*F* _(2, 21)_ = 1.64, *p* = 0.22], Table S1). For the test data, The RMANOVA showed a significant effect for Context [*F* _(1, 21)_ = 26.45, *p* < 0.001, *η^2^G* = 0.28] but not for Treatment [*F* _(2, 21)_ = 0.07, *p* = 0.93] or Context * Treatment interaction [*F* _(2, 21)_ = 0.35, *p* = 0.71, Fig. 4a]. The mean Generalization Index were similar between all groups [*F* _(2, 21)_ = 0.29, *p* = 0.75, Fig. 4b] and this lack of effect was further confirmed by the likelihood-ratio chi-square test [χ^2^ _(2, 24)_ = 0.00, *p* = 1.00, Fig. 4c].

**Figure 4:**
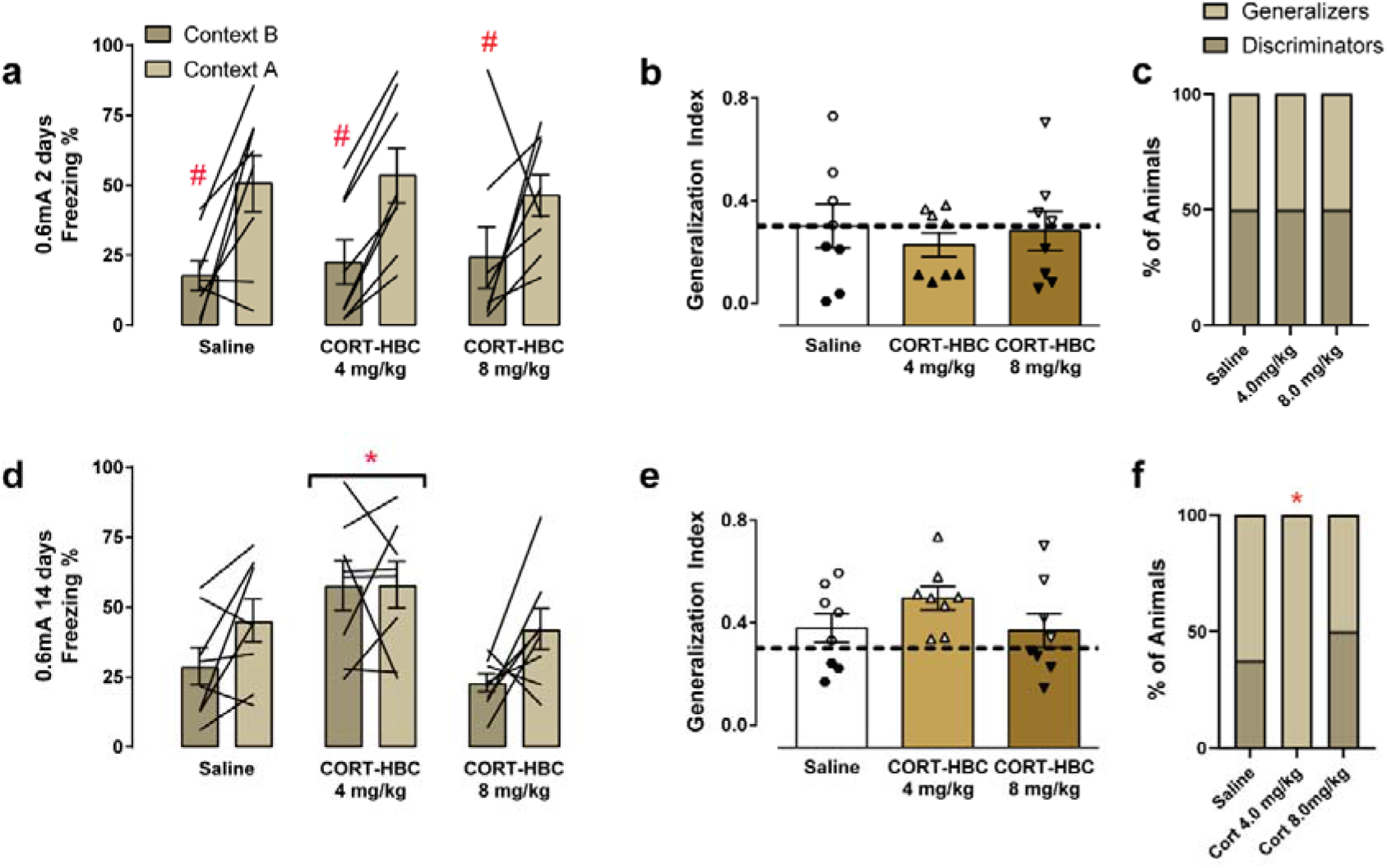
Corticosterone administration after moderate CFC training intensity increases contextual fear generalization at 14 days. (a, d) Mean freezing percentage ± Standard error of the mean (S.E.M) shown by rats exposed to the novel context (B) and then to the training context (A) (a) 2 or (d) 14 days after training. Lines represent individual freezing data for each rat in context B and then A. (#) p < 0.001 when compared to within-subject freezing expressed in Context A. (*) p = 0.01 when compared to the 8.0mg/kg and Saline groups. (b, e) Mean Generalization Index ± S.E.M. [B/(A+B)] (b) 2 or (d) 28 days after CFC. The dotted line represents the threshold between “discriminators” and “generalizers”. Filled symbols represent the animals considered discriminators and the symbols unfilled represent the ones considered generalizers. (c, f) Percentage of generalizer and discriminator animals per group after (c) 2 or (f) 14 days after CFC. (*) p < 0.05 in the proportion test. Number of rats for each group: 2 days, Saline (N = 8), CORT-HBC 4 mg/kg (N = 8), 8 mg/kg (N = 8). 14 days, Saline (N = 8), CORT-HBC 4 mg/kg (N = 8), 8 mg/kg (N = 8).

On the other hand, results from the second set of animals, tested 14 days after training, indicate that rats injected with 4.0 mg/kg of CORT-HBC showed increased freezing in Context B when compared to the other groups, while maintaining the same level of freezing in Context A (Fig. 4d). Freezing data from the training session did not show any difference between groups (Treatment [*F* _(2, 21)_ = 1.61, *p* = 0.22], Table S1). During the test session, freezing data from Context B did not show homogenous variance (*p* < 0.05); therefore, they were transformed using the square root function. The RMANOVA showed a significant effect for Context [*F* _(1, 21)_ = 6.83, *p* = 0.02, *η^2^G* = 0.11] and Treatment [*F* _(2, 21)_ = 5.27, *p* = 0.01, *η^2^G* = 0.24]. However, the interaction between Context and Treatment was not significant [*F* _(2, 21)_ = 1.63, *p* = 0.22]. Post hoc tests of the Treatment main effect showed that rats treated with 4.0 mg/kg of CORT-HBC showed higher overall freezing times than the ones treated with Saline (*t* _(21)_ = 2.54, *p* = 0.048, *d* = 1.11) or 8.0 mg/kg of CORT-HBC (*t* _(21)_ = 3.02, *p* = 0.02, *d* = 1.32). Overall freezing times of Saline and 8.0 mg/kg-CORT-HBC groups were not statistically different from each other (*t* _(21)_ = −0.48, *p* = 0.88; Fig. 4d). As there was no interaction effect, it is not possible to perform pos-hoc tests to compare the freezing behavior in contexts A and B of each group, and the significant effect for Context indicates that all animals spent in average more time in freezing in the training context than in the novel context. To further assess between-group differences, additional Bonferroni corrected one-way ANOVAs were used to compare freezing in Context A or B, separately (Table 1 – 0.6mA / 14 days). These ANOVAs showed no significant effect between groups tested in the training context (p = 0.36), but a significant effect between groups tested in the novel context [*F* _(2, 21)_ = 7.23, *p* < 0.01, *η^2^p* = 0.41]. The post-hoc analyses confirmed that, in tests performed in context B, rats in the 4.0 mg/kg-CORT-HBC group showed higher freezing times when compared to Saline (*p* = 0.02) and 8.0 mg/kg-CORT-HBC (*p* <0.01) groups, while these two groups showed no difference between them (*p* = 0.86). Moreover, although the comparison between generalization indices did not show any differences [*F* _(2, 21)_ = 1.58, *p* = 0.23, Fig. 4e], the likelihood-ratio chi-square test showed a significant effect in the distribution of generalizers in the 4.0mg/kg-CORT-HBC group (N = 8) when compared the 8.0mg/kg-CORT-HBC (N = 4) and Saline groups [N = 5, *χ^2^* _(2, 24)_ = 7.30, *p* = 0.03, Fig. 4f].

In summary, according to two of the pre-established measures (increased freezing levels in Context B and a higher proportion of generalizing animals), we may infer that treatment with CORT-HBC at the dose of 4.0 mg/kg, but not 8.0 mg/kg, immediately after the moderate CFC training, enhanced fear memory generalization 14 days later, without interfering in recent memory accuracy.

## DISCUSSION

The results reported here indicate that CORT-HBC modulates contextual fear memory specificity differently for mild and moderate training intensities. Our findings in the Saline groups replicate previous findings that mild CFC (3x 0.3mA) elicit similar levels of freezing during recent retrieval tests but discriminate between contexts at remote timepoints (dos Santos Corrêa et al., 2019). Contrary to our hypothesis, the treatment with CORT-HBC after this training intensity promoted a contextual memory accuracy in the recent (2 days) retrieval test, and increased the discriminative fear response of animals at the 28^th^ day, compared to the control rats (even though only one of the variables measured was significant and only with the higher dose). Additionally, on the 28^th^ day retrieval test, the 4.0 mg/kg dose seemed to lower overall freezing levels while still maintaining contextual discrimination. It is possible that this dose of CORT-HBC, given immediately after a mild CFC training, is impairing the strength or the persistence of the contextual memory, without interfering in its specificity (according to the index of generalization and the proportion of “generalizer” animals). Curiously, freezing levels of this group (4 mg/kg-CORT-HBC) at the 28th day was very similar to that observed in all groups at the 42 day time point. Therefore, another possibility is that post-training injections of 4 mg/kg of CORT-HBC, immediately after a mild CFC training, is anticipating the freezing levels that would be expected at more remote time points.

Dos Santos Corrêa and colleagues (2019) also showed that rats trained with the moderate CFC intensity (3x 0.6mA) showed contextual discrimination in the recent retrieval tests (2 days after training) and then generalized fear between the training context and a new context at the 28^th^ day after training. It is important to note that another group of rats tested 14 days after training showed a generalized memory in the first experiment. However, this result was not replicated in a second experiment reported in the same study in which another group of rats trained under the same parameters discriminated across contexts at this intermediate time point (dos Santos Corrêa et al., 2019). We did additional *post hoc* analysis of that data and the results thereof revealed that the replication group had lower mean CORT levels when compared to the rats from the original experiment (Fig S1, supplementary material) which raised the question whether this moderate training intensity and time point of testing combined are more susceptible to CORT’s modulation on memory accuracy. In the present study, we hypothesized that animals trained with this footshock intensity and treated with CORT-HBC would show a facilitation to generalize contextual fear. Our results indicate that treatment with CORT-HBC at the dose of 4.0 mg/kg, but not 8.0 mg/Kg, increased generalized freezing to the novel context, without increasing freezing in the training context, 14 days after the moderate CFC training. These data suggest that treatment with CORT-HBC may ‘tip the balance’ towards fear generalization at the 14^th^ day after training following an inverted U-curve dose-response function, with the dose of 4.0 mg/kg eliciting fear generalization and the dose of 8.0 mg/kg maintaining some contextual discrimination similarly to the saline group. This result suggests that there is an optimal glucocorticoid activation after a highly arousing experience that modulates the consolidation of generalized memory traces, which agrees with previous data reported in rats (Pedraza et al., 2016; Roozendaal and Mirone, 2020). However, it is important to emphasize that all comparisons across timepoints are done at the group level because different sets of animals were used for each time interval and footshock intensity, therefore this inference is only correlational.

Our results indicate, therefore, that acute post-training CORT treatment elicits different behavioral outcomes, contingent on the CFC training intensity. Likewise, studies in humans suggest that stress may enhance memory accuracy (Kensinger et al., 2007; Segal et al., 2012), but there are divergent results on the literature showing an increase of false content in episodic memories and reduced accuracy after learning under stress (Rimmele et al., 2011; Schwabe et al., 2009; Van Ast et al., 2014). We emphasized, however, that our study used a post-training CORT treatment that is not directly translatable to some of the studies in humans cited above, since their emotional or pharmacological interventions were performed at different phases of memory encoding, not only after training. Regardless of the intervention’s timepoint, these studies indicate that stress at the time of encoding can act either enhancing or impairing memory accuracy. As supporting evidence in rodents to our results, that suggest that acute CORT treatment immediately after a highly arousing CFC training temporally modulates contextual specificity, Pedraza and colleagues reported that pre-training CORT inhibition by metyrapone maintained the accuracy of contextual fear memory up to 25 days after a strong CFC training in rats, in contrast to vehicle treated rats that showed generalized fear at this timepoint. Roozendaal and Mirone (2020) showed that post-training CORT treatment induced a generalized strengthening of memory in a modified version of the inhibitory avoidance task during recent retrieval, while noradrenergic activation after training induced an increase in the accuracy of the association between context and shock, which indicates that both hormones act synergistically to consolidate contextual features of an aversive event. These findings together point to an important role played by the level of arousal during acquisition on CORT’s modulation of contextual fear memories. It is important to note that the latter results diverge slightly from our results with mild training intensity, in which CORT treatment seems to enhance memory specificity up to 28 days, but it can be rationalized by differences in rat lineage, quantity and intensity of footshocks received, as well as by the different behavioral outcomes quantified during the test and the different intrinsic features of the CFC and inhibitory avoidance tasks. On the other hand, our results with the moderate CFC training agree with this hypothesis that CORT increases fear generalization. More studies investigating the effects of blocking the glucocorticoid and noradrenergic systems are needed to better understand the role of these hormones in the specificity of contextual fear memory over time.

Additional evidences in humans showed that increasing stress before training elicited reorientation of attention toward salient contextual features (Clewett and Murty, 2019; Schwabe, 2017). These evidences suggest that more arousing experiences elicit increased noradrenergic activity (Roozendaal and Hermans, 2017) that, followed by appropriate CORT expression, elicit optimal consolidation of the memory trace. Therefore, our results reported here can be explained by the increasing levels of arousal during mild or moderate CFC sessions that interact with circulating CORT in response to the aversive situation, prompting differential processes of systems consolidation (Bahtiyar et al., 2020; dos Santos Corrêa et al., 2019). Mild intensity CFC may induce a process of systems consolidation in which exogenous CORT sustains an increased memory specificity at remote timepoints, whereas higher intensity CFC possibly recruits brain regions associated with more rigid learning, such as the striatum, and induces a different process of systems consolidation in which exogenous CORT facilitates time-dependent fear generalization.

Our data raise the question of how CORT would prompt different routes of systems consolidation after low or highly arousing events. It is possible to conjecture that memory acquisition under stress shifts the focus of attention to central features of the aversive event and less on its contextual details (Christianson and Loftus, 1987). This shift of focus is associated with a shift of functional activity from areas involved in contextual consolidation, such as the hippocampus (Addis and Schacter, 2008), to cortical areas implicated in semantic representations, such as the parahippocampal cortex (Binder et al., 2009). The hippocampal memory system is thought to encode separate representation of events and allow the acquisition of flexible memory representations (Shohamy and Wagner, 2008). Rapid glucocorticoid activity would act in association with the noradrenergic system in the amygdala, prompting brain-wide recruitment of brain regions associated with semantic representations and the habit memory system, favoring the association of threat-related signs to the memory trace and faster incorporation of the trace into preexisting semantic schemas (Quaedflieg and Schwabe, 2018; Schwabe, 2017; Schwabe and Wolf, 2012). If stress induces a shift from cognitive to habit learning, which lacks this flexibility for consolidating contextual representations (Myers et al., 2003), it could also increase generalization, which agrees with studies done in humans (McGlade et al., 2019; Rimmele et al., 2011; Schwabe et al., 2009). In contrast, low arousing events would maintain memory dependence on the hippocampal system.

Our results agree with this supposition. In one hand, mild CFC possibly did not increase rapid noradrenergic and glucocorticoid activation to levels at which there would be a shift of memory systems towards striatal learning (Quirarte et al., 1998; Roozendaal and Hermans, 2017), maintaining memory accuracy up to 42 days. Additional treatment with exogenous CORT further affected memory specificity, increasing contextual discrimination up to 28 days, as shown by the rats trained with 0.3mA footshocks and treated with CORT-HBC. On the other hand, moderate CFC possibly reached the necessary stress levels to induce the shift of memory systems towards reflexive learning. Exogenous CORT for this training intensity acted facilitating the consolidation of a rigid and contextually poor memory trace (Vogel et al., 2016) that became generalized over time.

Surprisingly, we did not find an enhancement in fear memory strength after post-training treatment with CORT when rats were exposed to the training context, as it was previously reported (Abrari et al., 2009; Cordero and Sandi, 1998; Kaouane et al., 2012; Pugh et al., 1997; Roozendaal and Mirone, 2020). Even though there are other studies that also did not report this enhancement after treatment with CORT (Bueno et al., 2017; Marchand et al., 2007), we hypothesize that the lack of effect in our experiment is due to differences in training parameters, experimental design or task-related features. It is possible that the order of context exposure during test sessions in the present study, i.e., first in the novel context and then in the conditioning context, may have prevented the observation of the CORT-induced enhancement in contextual memory strength. Some of the studies that have investigated stress hormones effects on episodic-like memory strength and accuracy used a counterbalanced exposure to different contexts in the inhibitory avoidance task and reported no order effect (Atucha and Roozendaal, 2015; Roozendaal and Mirone, 2020). However, pilot experiments done in our laboratory using a counterbalanced testing after a moderate CFC training (3 footshocks of 0.6 mA) showed an order effect: animals tested first in the training context and then in the novel context showed an accurate memory either 2 or 28 days after training, whereas rats tested first in the novel context showed an accurate memory only 2 days after training, and a generalized memory at the 28th day (unpublished results). These results suggest that, for our experimental parameters, the exposure to the conditioning context first, during testing sessions, influences remote memory generalization in the CFC task. As our main interest was to verify whether CORT administration would modulate memory generalization over time, we chose to test animals first in the new context.

In conclusion, this study strengthens the hypothesis that CORT modifies time-dependent memory generalization, associated with the process of systems consolidation. In the experimental design described here, post-training exogenous CORT after mild CFC seemed to increase recent and remote memory accuracy and to accelerate freezing levels expected only in more remote timepoints. On the other hand, CORT injections after moderate CFC seemed to facilitate time-dependent generalization. This differential modulation of CORT on fear memory specificity, contingent to training intensity, conveys strong support to the hypothesis of arousal-induced shift in memory systems during CFC acquisition and consolidation.

## Acknowledgements

We thank the valuable support from Vitor Farhat Fernandes, Lorena Vido Lopes and Gabriela Iramina Gomes for helping in some of the experiments. We also thank the work done by the technicians that manage the vivarium. The authors would like to thank the members of the Research group in Neurobiology of Learning and Memory at UFABC (MANAs) for scientific advice and discussion.

## Research data for this article

The data used, collected and analyzed for all results here presented can be found at ([dataset] Dos Santos Corrêa, M.; Fornari, 2020) http://dx.doi.org/10.17632/7xvshh2s59.1

## Author Statement

MdSC helped conceptualizing, curating data, doing formal analysis and experiments, planning methodology and writing the original draft. BdSV and BSM helped doing experiments and reviewing the manuscript. PAT and TLF helped reviewing the manuscript. RVF helped conceptualizing, acquiring funds, planning methodology, administrating and supervising the project and reviewing the manuscript.

