## Supplementary material for "Corticosterone differentially modulates time-dependent fear generalization following mild or moderate fear conditioning training in rats": Suplemental Material (FigS1 and Tables S1 and S2)

**Figure S1.**

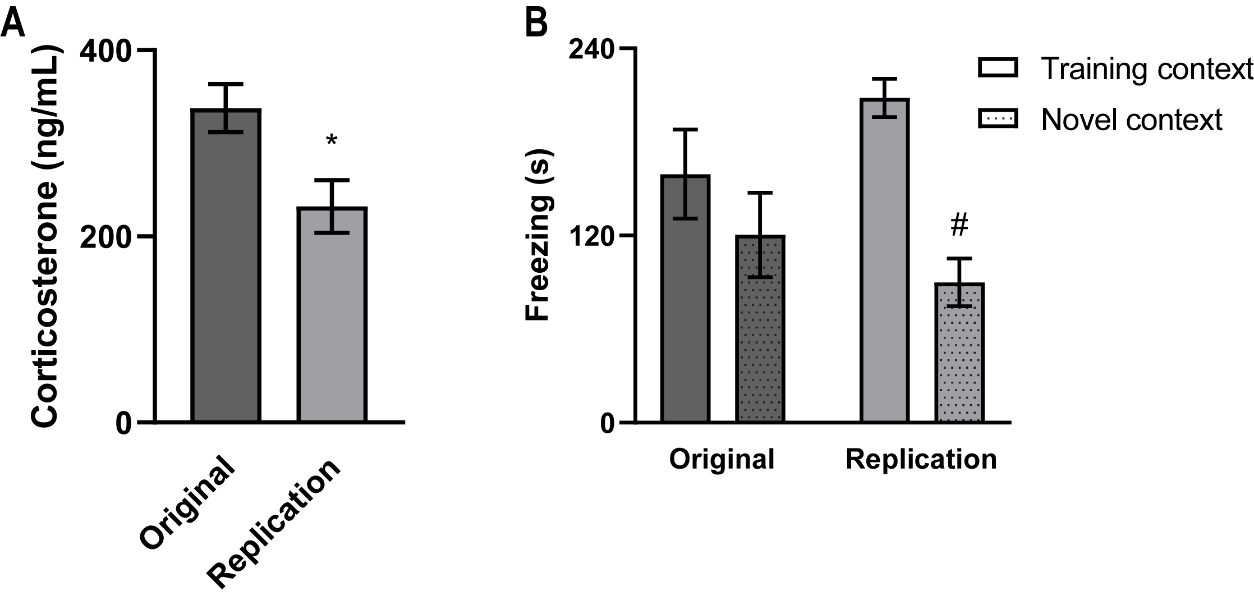

Comparison of two experiments (original x replication) carried out with different animal cohorts that were trained with 3 x 0.6 mA shocks and tested in the same training context or in a novel context after 14 days. The data were extracted from a previously published dataset [1] and re-analyzed here.

(a) Mean post-training Corticosterone levels (ng/mL) for animals from the original and replication experiments. Data was compared using a Welch’s t-test, *t* (37.5) =2.76, *p*=0.009. N = 16 for the original and 24 for the replication groups.
(b) Mean freezing times (s) of animals trained with 3 footshocks of **0.6mA** and tested after **14 days** either in the **training or the novel context**, in the **original and replication** experiments. Different animals were used for each of the four groups. Animals from the original experiment showed similar freezing times in both contexts, whereas in the replication experiment, rats tested in the training context presented higher freezing levels than those tested in the novel context. Data was compared using multiple t-tests with the Hold-Sidak correction method. Original experiment: *t* (14) =0.99, adjusted p=0.34; Replication experiment: *t* (22) =6.040, adj. p<0.001. N = 8 for the training and 8 for the novel contexts in the original groups, N=12 for the training and 12 for the novel contexts in the replication groups.

**Methods**

Data used here is published at <https://data.mendeley.com/datasets/n5zdf72x6s/5>. The statistical analysis shown above were done after the publication of a previous article and are considered novel results.

*Subjects:*

Three-month old male Wistar rats, originating from *Instituto Nacional de Farmacologia* (INFAR-UNIFESP; (total n = 40; weighing between 275-385g at time of training) were kept in controlled conditions of temperature (23 ± 2°C) and light/darkness cycle of 12:12 hours (light phase starting at 7am). All procedures were conducted according to the guidelines and standards of CONCEA - *Conselho Nacional de Controle de Experimentação Animal* (Brazilian Council of Animal Experimentation) and were previously approved by the Ethics Committee on Animal Use - UFABC (CEUA - protocol number 7479070916).

*Experimental design and ELISA*

All procedures are thoroughly described previously [2].

**Table S1**

| **Table S1: One-way ANOVAs for the freezing times in the last minute of the training session** | | | | | | |
| --- | --- | --- | --- | --- | --- | --- |
| **Experiment** | **Post-shock freezing % ± S.E.M*** | | | **F** | **df** | **p-value** |
|  | Saline | CORT-HBC 4.0 mg/kg | CORT-HBC 8.0 mg/kg |  |  |  |
| 0.3 mA\|2 days | 45.3 ± 10.4 (7) | 48.0 ± 8.93 (8) | 42.1 ± 11.5 (7) | 0.09 | 2,19 | 0.92 |
| 0.3 mA\|28 days | 23.3 ± 5.32 (8) | 28.9 ± 5.80 (8) | 31.1 ± 6.02 (8) | 0.50 | 2,21 | 0.61 |
| 0.3 mA\|42 days | 36.9 ± 9.12 (8) | 37.9 ± 7.09 (7) | 45.8 ± 11.3 (8) | 0.27 | 2,20 | 0.77 |
| 0.6 mA\|2 days | 44.4 ± 9.38 (8) | 26.5 ± 5.77 (8) | 43.0 ± 10.4 (8) | 1.64 | 2,21 | 0.22 |
| 0.6 mA\|14 days | 28.2 ± 10.0 (8) | 41.4 ± 9.33 (8) | 42.3 ± 8.75 (8) | 1.61 | 2,21 | 0.22 |
| **In parenthesis, the number of rats in that group* | | |  |  |  |  |

**Table S2**

| **Table S2: Bonferroni corrected One-way ANOVAs for the freezing times in the training (A) or novel (B) contexts and *post hoc* tests** | | | | | | |
| --- | --- | --- | --- | --- | --- | --- |
| **Dosage selection** | | | | | | |
| **ANOVAs** | | **F** | **df1** | **df2** | **p** | **η²p** |
| Freezing B |  | 1.03 | 6 | 40 | 0.42 | 0.13 |
| Freezing A |  | 0.53 | 6 | 40 | 0.78 | 0.07 |
| **0.3mA\|2 days** | | | | | | |
| **ANOVAs** | | **F** | **df1** | **df2** | **p** | **η²p** |
| Freezing B |  | 1.44 | 2 | 19 | 0.26 | 0.13 |
| Freezing A |  | 1.16 | 2 | 19 | 0.34 | 0.11 |
| **0.3mA\|28 days** | | | | | | |
| **ANOVAs** | | **F** | **df1** | **df2** | **p** | **η²p** |
| Freezing B |  | 7.23 | 2 | 21 | **<0.01*** | 0.41 |
| Freezing A |  | 5.60 | 2 | 21 | **0.01*** | 0.35 |
| **Post hoc tests Freezing A** | | **Mdiff** | **t** | **df** | **p** | **d** |
| Saline (FreezA) | CORT-HBC 4.0 (Freez A) | 4.71 | 3.17 | 21 | **0.01*** | 1.58 |
| Saline (FreezA) | CORT-HBC 8.0 (Freez A) | 0.97 | 0.65 | 21 | 0.79 | 0.33 |
| CORT-HBC 4.0 (Freez A) | CORT-HBC 8.0 (Freez A) | -3.74 | -2.52 | 21 | **0.05*** | -1.26 |
| **Post hoc tests Freezing B** | | **Mdiff** | **t** | **df** | **p** | **d** |
| Saline (FreezB) | CORT-HBC 4.0 (Freez B) | 3.08 | 2.55 | 21 | **0.047*** | 1.28 |
| Saline (FreezB) | CORT-HBC 8.0 (Freez B) | 2.97 | 2.46 | 21 | 0.06 | 1.23 |
| CORT-HBC 4.0 (Freez B) | CORT-HBC 8.0 (Freez B) | -0.11 | -0.09 | 21 | 1.00 | -0.04 |
| **0.3mA\|42 days** | | | | | | |
| **ANOVAs** | | **F** | **df1** | **df2** | **p** | **η²p** |
| Freezing B |  | 0.68 | 2 | 20 | 0.51 | 0.06 |
| Freezing A |  | 0.12 | 2 | 20 | 0.89 | 0.01 |
| **0.6mA\|2 days** | | | | | | |
| **ANOVAs** | | **F** | **df1** | **df2** | **p** | **η²p** |
| Freezing B |  | 0.16 | 2 | 21 | 0.85 | 0.02 |
| Freezing A |  | 0.16 | 2 | 21 | 0.86 | 0.02 |
| **0.6mA\|14 days** | | | | | | |
| **ANOVAs** | | **F** | **df1** | **df2** | **p** | **η²p** |
| Freezing B |  | 7.23 | 2 | 21 | **<0.01*** | 0.41 |
| Freezing A |  | 1.07 | 2 | 21 | 0.36 | 0.09 |
| **Post hoc tests Freezing B** | | **Mdiff** | **t** | **df** | **p** | **d** |
| Saline (FreezB) | CORT-HBC 4.0 (Freez B) | -3.64 | -3.0 | 21 | **0.02*** | -1.5 |
| Saline (FreezB) | CORT-HBC 8.0 (Freez B) | 0.64 | 0.52 | 21 | 0.86 | 0.26 |
| CORT-HBC 4.0 (Freez B) | CORT-HBC 8.0 (Freez B) | -4.3 | -3.5 | 21 | **<0.01*** | -1.76 |
| Bonferroni corrected ANOVAs significance is 0.025. | | | | | | |

References

[1]	M. [dataset] dos Santos Corrêa, V. Barbara dos Santos, G. Gabriel David Vieira, de P. Joselisa Péres Queiroz, T. Paula Ayako, and R. V. Fornari, *Relationship between footshock intensity, post-training corticosterone release and contextual fear memory specificity over time (raw data)*. v5: Mendeley Data, 2019.

[2]	M. dos Santos Corrêa, B. dos S. Vaz, G. D. V. Grisanti, J. P. Q. de Paiva, P. A. Tiba, and R. V. Fornari, “Relationship between footshock intensity, post-training corticosterone release and contextual fear memory specificity over time,” *Psychoneuroendocrinology*, vol. 110, no. September, p. 104447, Dec. 2019, doi: 10.1016/j.psyneuen.2019.104447.
